# Sensing and seeing associated with overlapping occipitoparietal activation in simultaneous EEG-fMRI

**DOI:** 10.1101/2020.07.08.193326

**Authors:** Catriona L. Scrivener, Asad Malik, Michael Lindner, Etienne B. Roesch

## Abstract

The presence of a change in a visual scene can influence brain activity and behaviour, even in the absence of full conscious report. It may be possible for us to *sense* that such a change has occurred, even if we cannot specify exactly where or what it was. Despite existing evidence from electroencephalogram (EEG) and eye-tracking data, it is still unclear how this partial level of awareness relates to fMRI BOLD activation. Using EEG, functional magnetic resonance imaging (fMRI), and a change blindness paradigm, we found multi-modal evidence to suggest that *sensing* a change is distinguishable from being *blind* to it. Specifically, trials during which participants could detect the presence of a colour change but not identify the location of the change (*sense* trials), were compared to those where participants could both detect and localise the change (*localise* or *see* trials), as well as change blind trials. In EEG, late parietal positivity and N2 amplitudes were larger for *localised* changes only, when compared to change blindness. However, ERP-informed fMRI analysis found no voxels with activation that significantly co-varied with fluctuations in single-trial late positivity amplitudes. In fMRI, a range of visual (BA17,18), parietal (BA7,40), and midbrain (anterior cingulate, BA24) areas showed increased fMRI BOLD activation when a change was *sensed*, compared to change blindness. These visual and parietal areas are commonly implicated as the storage sites of visual working memory, and we therefore argue that sensing may not be explained by a lack of stored representation of the visual display. Both *seeing* and *sensing* a change were associated with an overlapping occipitoparietal network of activation when compared to *blind* trials, suggesting that the quality of the visual representation, rather than the lack of one, may result in partial awareness during the change blindness paradigm.

## Introduction

It is common for us to overestimate the amount of information that we can process and store about the world around us. Although we may assume that we would notice a cyclist entering the path of our car, or if a building on our street changed in colour, we very often miss these occurrences (Simons, 2000; Simons & Levin, 1997). The failure to detect changes between visual scenes is known as change blindness, and is used as evidence to suggest that our internal representation of the outside world is not as complete as once thought (Rensink, 2004; Noe et al., 2000). When changes to an image are disrupted in some way, for example by a distractor image or a visual saccade, we cannot use visual transients (or motion) to detect them, and are often blind to the difference (Rensink et al., 1997; Kanai & Verstraten, 2004).

It was previously assumed that if we are blind to a change then we cannot provide any information about it, and that the change should not influence our behaviour in any way. Blindness to changes is thought to result from a lack of detailed representation about the pre- and post-change scenes, or an inability to successfully compare the two (Simons, 2000). If this is the case, then our knowledge when we are blind to changes should be equivalent to that when there is no change at all. Anecdotally, this does not align with the experience of observers in a change blindness experiment; it is common for them to remark that they suspected something had changed, but that they were not sure about its nature or location. This experience appears to be phenomenologically different from complete change blindness, but how this difference is reflected in behavioural and neuroimaging data is unclear.

In the domain of visual consciousness, there is a recurring debate on the nature of visual awareness; whether it is graded, or dichotomous, or a combination determined by the context. In the Global Neuronal Workspace Theory (Dehaene & Naccache, 2001; Dehaene et al., 2006), it is posited that awareness arises when inputs cross a threshold for ‘ignition’, resulting in the distribution and maintenance of information within a ‘global workspace’. Based on this proposal, conscious awareness is a dichotomous state, as only inputs selected by attention can spark the activation of the global workspace. This consists of a large network of connected regions, including prefrontal and parietal regions as well as the thalamic nuclei and basal ganglia (Dehaene & Changeux, 2011). Therefore, conscious awareness requires directed attention and activation of a distributed frontal-parietal network, in an ‘all-or-nothing’ fashion.

In accordance with this, fMRI studies specifically investigating change blindness report that detected changes are associated with greater activation in the parietal lobe, dorsolateral prefrontal cortex, and fusiform gyrus, when compared to changes that are missed (Beck et al., 2001). Further, detected changes compared with correctly categorised no-change trials revealed activation in a wider network including the inferior, superior, and medial temporal gyrus, anterior interparietal sulcus, precuneus, central sulcus, inferior frontal gyrus, anterior cingulate cortex, putamen, pulvinar, and cerebellum (Pessoa, 2004). A similar pattern was identified for false alarm trials, where participants reported a change when no change occurred, suggesting that activity was related to the participants’ perception of the change rather than properties of the visual stimulus. Overall, few regions were specifically activated when participants exhibited change blindness.

However, this ‘all-or-nothing’ explanation of visual awareness does not align with our subjective experience of the world. Based on participants’ report of a sense for something changing, we might conclude that awareness is graded. This allows for a level, or levels, of awareness lying somewhere on a continuum between full and absent awareness. In an early experiment, Rensink (2004) suggested the presence of a *sense* condition, in which observers could detect a change without fully identifying it. Observers were asked to indicate when they ‘thought’ that something had changed, and then again when they were certain of it. He argued that this *sense* condition is both phenomenologically and perceptually distinct from the traditionally reported *see* condition in which participants are fully aware of what change occurred.

This definition has been extended and explored using electroencephalogram (EEG) and eye-tracking, with a range of results suggesting a richer visual experience than either “yes I saw a change” or “no I didn’t see anything” (Busch et al., 2009; Fernandez-Duque & Thornton, 2003; Kimura et al., 2008; Lyyra et al., 2012; Thornton & Fernandez-Duque, 2001; Howe & Webb, 2014; Chetverikov et al., 2018; Reynolds & Withers, 2015; Lyyra et al., 2012; Galpin et al., 2008). The distinction could be described by the ‘partial awareness hypothesis’ (Kouider et al., 2010; Kouider & Dehaene, 2007). While the mechanism of awareness can still be considered dichotomous and dependent on an ignition threshold, the level of detail contained within the workspace is variable. Stimuli can be represented with varying detail, based on factors such as stimulus strength, therefore giving rise to graded knowledge of its contents.

In a previous EEG experiment (Scrivener et al., 2019) we distinguished between trials in which participants could detect the presence of a colour change but not identify the location of the change (*sense* trials), versus those where participants could both detect and localise the change (*localise* trials). We chose to measure several ERPs that are commonly linked to visual attention and awareness, including the visual P1 and N1, visual awareness negativity (VAN), N2pc, and late positivity (LP) (Koivisto & Revonsuo, 2010; Förster et al., 2020). Although suggested as one of the earliest reflection of conscious visual awareness (around 200 ms after stimulus onset), we found no statistically significant differences in the VAN ERP across conditions, contrary to previous findings (Förster et al., 2020; Koivisto et al., 2008; Wilenius & Revonsuo, 2007; Busch et al., 2009).

In a similar time window, the N2pc is characterised by an increased negativity at visual electrodes contralateral to the change location, and is increased for aware versus unaware trials (Schankin & Wascher, 2007; Luck & Hillyard, 1994). In our previous results, both awareness conditions (*localise* and *sense*) were significantly different to trials with no change detection (*blind* trials), suggesting that the N2pc is not dependent on explicit awareness. It is possible that *sense* trials elicited a shift in attention to the correct hemifield of change (and therefore an N2pc was detected), but that this was not specific enough to determine the exact location of the change.

Within the late positivity range (400 - 600 ms after change onset), all conditions were significantly different from one another. The LP overlaps with the P3 component at central parietal electrode sites, and is often associated with conscious aspects of task processing (Koivisto et al., 2009; Busch et al., 2009; Railo et al., 2011). Overall, it appears that simply ‘detecting’ a change can be distinguished from ‘describing’ a change, in both subjective and neuroimgaging results, and that participants can *sense* a change without complete knowledge of what occurred.

The main aim of this experiment was to examine the existence and nature of the *sense* condition in the change blindness paradigm, using combined EEG-fMRI and behavioural measures. While a range of evidence posits a distinction between *sense* and *blind* conditions in EEG data, no such distinction has been made for the *sense* condition in change blindness using fMRI. One criticism of the sensing hypothesis is that participants who *sense* a change are simply applying a more liberal response criterion when completing the task, and in fact are not aware of the change at all (Simons et al., 2005). Similarly, implicit awareness of changes could also be explained by explicit mechanisms such as guessing or a process of elimination (Mitroff et al., 2002). If this is the case, then we would expect to find no significant differences between *sense* and *blind* trials in fMRI BOLD activation. This result could also support the hypothesis of visual consciousness as dichotomous. However, if sensing lies somewhere on a continuum between aware and unaware, perhaps explained by varying precision of the stimuli representation within the global workspace, then BOLD activation for *sense* trials may be separable to both fully aware and change *blind* trials.

Further, we aimed to improve the respective temporal and spatial resolution of EEG and fMRI by measuring them simultaneously. In an extension to our previous EEG results (Scrivener et al., 2019), we investigated how EEG correlates of visual awareness relate to changes in fMRI BOLD. We therefore aimed to identify brain regions with BOLD activity that co-varied with activity in the EEG data, to detect possible sources or networks associated with awareness of changes.

## Materials and Methods

All materials and analysis methods were pre-registered in an open document on the Open Science Framework, where the data and analysis for this project can also be found (https://doi.org/10.17605/OSF.IO/W6BH3). Structural images were defaced using Brainstorm3 (Tadel et al., 2011) in MATLAB (MathWorks, Inc., version 2014a) with SPM8.

### Participants

Twenty one right-handed subjects (mean ± SD, age = 21 ± 3.6, 6 male) with no history of psychiatric or neurological disorders participated in this EEG-fMRI study. All had corrected-to-normal vision and were not colour blind (based on self report). The experiment was approved by the University of Reading ethics committee (UREC: 16/120), and was conducted in accordance with the Declaration of Helsinki (as of 2008). All participants gave informed consent to take part, including consent to share their anonymised data. For EEG and behavioural analysis, one participant was removed due to failure to remove MRI related artifacts from the EEG, leaving N=20. Four additional participants were removed from the fMRI and EEG-fMRI analysis for having motion greater than one voxel size in the fMRI data, leaving N=16.

### Stimuli and procedure

A change blindness task was presented using Psychtoolbox (Kleiner et al., 2007), on a 1920 × 1080 LCD monitor with a 60 Hz refresh rate. The paradigm was displayed on a screen displayed approximately 47cm away from the centre of the scanner bore. This was viewed by the participant through a mirror mounted onto the coil, at approximately 12cm from the participant’s eyes. In their left hand, the participant held an alarm ball, and in their right they held a 4 key button box. They had to use all of the 4 keys to respond to the task. Participants were asked to fixate on a central fixation cross and identify changes between consecutive displays of coloured squares. These were interrupted by a short fixation display to facilitate the change blindness phenomenon (see figure 1 for details on display duration). On change trials, one of the squares changed colour from the first to the second display. On no-change trials, the displays were identical. This was followed by two or three questions, depending on the participant’s response to the first question.

**Figure 1.**
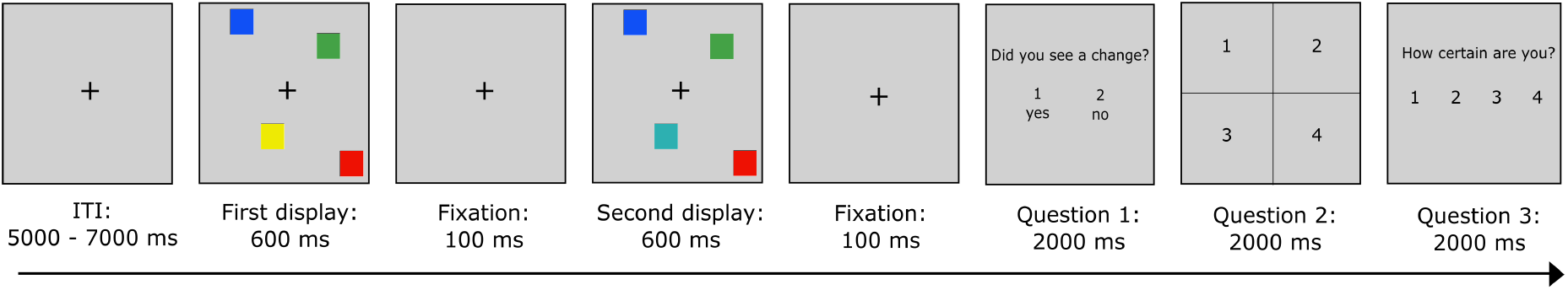
Illustration of the experimental paradigm. The number of squares presented varied from 2 to a maximum of 36. Question 1 asked ‘Did you see a change?’ to which participants could respond ‘Yes’ or ‘No’. Question 2 asked participants to localise the change, based on a grid from top left to bottom right. Question 3 asked how certain participants were of their responses, ranging from ‘1: Very Uncertain’ to ‘4: Very Certain’. If participants responded ‘no change’ to question 1, they were asked to press any button instead of the localisation response.

Question 1 asked ‘Did you see a change?’ to which participants could respond ‘yes’ or ‘no’. Question 2 asked participants to localise the change, based on a 2×2 grid from top left to bottom right. Question 3 asked how certain participants were of their responses, ranging from ‘1: Very Uncertain’ to ‘4: Very Certain’. If participants responded ‘no’ change to question 1, they were asked to press any button to ensure the same number of button presses were made during each trial. We did not ask participants who did not see the change to guess a location, as our hypotheses did not relate to ‘implicit’ change detection, as reported in Fernandez-Duque & Thornton (2000). Participants were asked to respond within a limit of two seconds for each question, and trials with any response missing were not included in further analysis.

This study had a within-subjects repeated measures design, and each participant completed 5 blocks of 50 trials, meaning a total of 250 trials. Of these 250 trials, 165 contained a change in coloured square, and the remaining trials contain no change. The ratio was not kept at 50/50, as the trials containing the change were of most interest for analysis. However, after the experiment participants were asked to report the percentage of trials that they believed contained a change. After each block of 50 trials, the participants were presented with a break screen, advising them to take a break. The participant was able to continue the experiment at their discretion by pressing any button on the button box. Before beginning the main task, participants were given a short block of 10 trials in which to practice responding to the paradigm with the button box. The data from this practice block was not analysed.

Difficulty was modulated in real time by adding and removing two squares from the display, based on the assumption that more distractors increases task difficulty (Vogel et al., 2005). This was to prevent floor and ceiling performance during the task as a result of individual differences (Luck & Vogel, 2013), and optimise for performance rather than to establish specific individual thresholds. Performance over the previous two trials was used to update the current trial; two consecutive correct answers added two squares, two incorrect deducted two squares, and one correct and one incorrect resulted in no change. The decision to increase or decrease the number of squares was made using responses to the localisation question (Q 2), as we were specifically interested in controlling the number of *sense* and *localise* trials. The number of squares always changed by two, to balance the number on the left right hemifields of the screen. The location of the change on each trial was random, but the change occurred an equal number of times on the left and right hemifield of the screen. The display was divided into 36 even sections, with 6 in each quadrant, within which the squares could appear. As the colour of the squares was not related to our main hypotheses, we used seven default MATLAB colours; blue, cyan, yellow, green, white, red, and magenta (MathWorks, Inc., version 2016b).

### Behavioural Analysis

The trials in which a change occurred were divided into three conditions: *blind* (no change detection), *localise* (change detection and localisation), and *sense* (change detection without localisation). Trials in which no change occurred were divided into *correct rejection* (no change reported) and *false alarm* (change incorrectly reported).

The number of false alarm trials was low, with a mean of 10 trials (range = 1 − 28, *SD* = 7.34), and therefore EEG analysis comparing *false alarm* to *sense* trials was not performed due to a lack of power. The percentage of *false alarm* trials was calculated in relation to the the total number of no-change trials, whereas the percentage of *sense* trials was calculated in relation to the total number of change trials.

Detection accuracy for each participant was calculated based on the percentage of change trials in which they correctly detected a change. Localisation accuracy was calculated as the percentage of correctly detected changes where the localisation was also correct. We also recorded each participant’s mean and maximum difficulty scores, with the maximum referring to the highest number of squares that were displayed to them during the experiment. Behavioural analysis was completed in JASP 2018 (version 0.8.2.0).

D’prime was calculated as a measure of participant response bias. This was calculated using the equation *d* = *z*(hit rate)−*z*(false alarm rate) (Stanislaw & Todorov, 1999). Response bias, or criterion, was also calculated, where *c* = −0.5 ∗ (*z*(hit rate) + *z*(false alarm rate)) (Stanislaw & Todorov, 1999). *c* = 0 indicates no response bias to either ‘yes’ or ‘no’ responses. *c* > 0 indicates a bias towards ‘no’ responses, with fewer hits and fewer false alarms. *c* < 0 indicates bias towards ‘yes’, with more hits but also more false alarms. We expected that participants would display a range of response strategies.

One problem faced in identifying a *sense* condition is that it is difficult to distinguish these trials from those where participants did not really see a change (similar to a false alarm during no change trials), or those where participants press the wrong response key (Simons & Ambinder, 2005; Mitroff et al., 2002). Rensink et al. (2004) found that reaction times when participants thought that they had seen a change were shorter for change trials than no-change trials, meaning that participants were slower when they were simply making a false alarm. Galpin et al. (2008) also found greater certainty associated with *sensing* during change trials, compared to *false alarms*. We therefore compared reaction times across awareness conditions, as well as between levels of confidence. As trial numbers were low, ‘very uncertain’ and ‘uncertain’ responses were combined, and ‘certain’ and ‘very certain’ were combined. Each awareness condition therefore had two levels of certainty; for example, *localise certain* and *localise uncertain*.

To establish if the location of the change influenced the likelihood that it was detected, we conducted two chi-square analyses. The first analysis divided the 6 × 6 grid of possible change locations into two conditions, outside and central. Changes occurring in any of the 20 outermost locations were considered to be outside changes, and the 16 central locations were considered to be central. We ran a 2 × 3 chi-square with the independent variables of location (outside/central) and awareness (*blind*/*localise*/*sense*), and the dependent variable as the frequency of trials within each condition, across participants. In the second analysis, we instead compared the side of the display in which the change occurred, resulting in a 2 × 3 chi-square for hemisphere (left/right) and awareness (*blind*/*localise*/*sense*).

### EEG data acquisition

EEG data was recorded with an MRI-compatible cap equipped with carbon-wired Ag/AgCL electrodes (Braincap MR) from 64 scalp positions according to the international 10-10 system. The reference electrode was placed at FCz and the ground at AFz. An additional ECG electrode was positioned on the back to measure heart rate. An MRI-compatible EEG amplifier was used (Brain-Amp MR, Brain Products) with a sampling rate of 5000Hz. This was positioned at the back of the scanner bore and connected using ribbon cables that were secured with sandbags. Impedance was kept below 10kΩ for EEG channels and 5kΩ for the ECG. EEG recordings were performed with Brain Vision Recorder Software (Brain Products) and timings kept constant using a Brain Products SyncBox to synchronise EEG with the MRI system clock.

### EEG pre-processing

Raw EEG data was pre-processed using Brain Vision Analyzer version 2.1 (Brain Products). Correction for the MR gradient artifact was performed using a baseline corrected sliding average of MR volumes (Allen et al., 2000). Removal of cardioballistic artifacts involved the subtraction of heartbeat artifacts on a second by second basis, using a sliding average of 21 (Allen et al., 1998). The delay was detected using the CBC detection solution, individually for each subject. Peaks were detected semi-automatically, with a manual check of the algorithm’s estimations. ICA (Infomax; Bell & Senjnowski, 1996) was then used to remove further BCG residual artifacts (range: 1 - 4 additional ICs removed per participant). As outlined in Debener (2005), the presence of visual P1 and N1 peaks in the averaged data after pre-processing was used as an indication of the successful removal of artifacts.

The data was downsampled to 500 Hz to reduce computation time and then filtered with a high-pass filter of 0.1 Hz to remove low frequency drift (Butterworth, 2nd order). A low-pass filter of 40 Hz and a notch filter of 50 Hz were chosen to remove line noise. Independent component analysis (ICA) was used to remove eye movement artifacts (Infomax; Bell & Senjnowski, 1996). Two components were removed for each participant; one corresponding to eye-blinks and the other to lateralised eye-movements.

Further analysis was completed using EEGLab (Delorme & Makeig, 2004). Trials were marked as outliers if any ERP value was greater than 3 standard deviations from the mean value of that ERP across all trials (using the MATLAB function ‘isoutlier’). Note that we only searched for outliers in the electrodes used for analysis (P07, P08, Cz, Pz, and CPz). Trials marked as containing outliers were excluded from further analysis (*M* = 7 trials, *SD* = 12.98), as well as those where a response to any question was not made within the response time (*M* = 2 trials, *SD* = 2.79).

Segments were then taken from −200 to 7000 ms to include the whole trial, and baseline corrected using a mean of the data within −200ms to 0ms, where 0ms was the start of the first display of coloured squares (see figure 1). We chose the baseline period to be before the first display onset, rather than the second, as we were interested in visual ERPs that occurred in response to the both displays. It has also been suggested that ERPs in response to the first presentation of stimuli are related to the subsequent perception of change (Pourtois et al., 2006).

### EEG Analysis

To identify the peaks of the visually evoked potentials (P1 and N1), a grand average ERP was calculated across all conditions and participants, as advised in Luck and Gaspelin (2017), from electrodes P07 and P08. From here, the peaks of interest were determined by identifying the local maxima/minima of the expected peaks, using the peak detection function in BrainVision Analyzer. The mean value within a window around the peak was used instead of the peak value, as the mean is more robust against noise (Luck, 2014). A window of 40ms around the mean was chosen as the appropriate window for visual ERPs P1 and N1. In relation to the first display onset, the first P1 was identified at 124ms, and the first N1 at 142ms. In relation to the second display onset, the second P1 was identified at 108ms, and the second N1 at 168ms.

Based on previous literature (Busch et al., 2010; Tseng et al., 2012; Fernandez-Duque et al., 2003), the N2pc was defined as the mean within 200-400 ms after the second display at occipital electrodes PO7 and PO8. Over central parietal electrodes Cz, CPz and Pz, the VAN was defined within a window of 130-330 ms after the second display, and the LP within a window of 400-600ms. We used window sizes of 200 ms, defined a-priori, in an attempt to be conservative given the large variation within the literature.

To assess how differences between early visual components across detection conditions were reflected at each stimulus presentation, P1 and N1 amplitudes were compared in two separate 2×3 repeated measures ANOVAs, with display (first/second) and awareness (*blind*/*localise*/*sense*) as the independent variables. Differences across hemispheres in the N2pc were analysed with another 2×3 repeated measures ANOVA, with the independent variables of hemisphere (contralateral/ipsilateral) and awareness (*blind*/*localise*/*sense*). Amplitudes of the VAN and the LP were compared in two separate repeated measures ANOVAs with awareness (*blind*/*localise*/*sense*) as the independent variable. Where Mauchly’s Test of Sphericity indicated that the assumption had been violated, Greenhouse-Geisser correction was used. All post-hoc comparisons were two-tailed, and corrected for multiple comparisons using false discovery rate where *q* = .05 (Benjamini & Hochberg, 1995). Effect sizes are reported as partial eta squared for ANOVA, and repeated measures Hedge’s g for t-tests (Lakens, 2013).

### Single-trial EEG Analysis

As listed a-priori in our pre-registration document on the OSF, we used two methods to extract the single-trial ERP values. The first method used the raw EEG time series, while the second used EEG values extracted from the ICA component with maximum correlation with our ERP of interest. Our reason for using both methods was to increase our sensitivity for extracting meaningful single trial values, given the reduced signal to noise ratio in EEG data recorded inside the MR environment.

#### Raw values

for each ERP time window, single-trial values were calculated as the mean amplitude within the predefined window for that peak. These values were then baseline corrected by subtracting the mean amplitude across the trial from which they were taken. Outliers were identified as trials where the amplitude was more than 3 standard deviations away from the mean amplitude for that ERP. As large artifacts can raise the mean amplitude, we added the additional classification of outliers at values +/−30 *µV* . These outlier values were replaced by the mean value across all other trials, as outlined in Bénar et al. (2007).

#### ICA derived values

this method was similar to that mentioned above, with the exception that the single-trial values were taken from a single ICA component, identified separately for each participant. First, ICA was computed on the pre-processed data for a single subject (FastICA in EEGLab; Hyvärinen & Oja, 1997). For each independent component (IC) extracted, a mean IC ERP was calculated by averaging the time course across all trials. The average IC ERP time courses were then correlated with the average ERP time course from the electrodes of interest in the pre-processed EEG data; for the LP this was the average ERP from the central electrodes (Cz, Pz, CPz). The IC component with the highest correlation with the ERP of interest was inspected to ensure that the topology was as expected; for the LP this was positivity over the central electrodes. Once selected, the single-trial values were extracted from the time series of this component, as described above. For some participants, the IC with the highest correlation was an artifact component, identified by visually inspecting the component’s time series, topography, and frequency spectrum in EEGLab. We also utilised the EEGLab function ‘ICLabel’ to aid classification of artifact components. When this was the case (3 participants), the IC with the next highest correlation was selected for that participant.

### fMRI recording

MRI data was acquired using a 3.0-T whole-body MRI scanner (Prisma, Siemens) and a 64 channel coil for functional imaging. Interleaved slices were recorded using a 2D echo planar imaging (EPI) sequence [repetition time (TR) 1630ms; echo time (TE) 30ms; flip angle 90ΰ; voxel size 3mm × 3mm; gap 3mm; encoding direction A to P; distance factor 20%; FOV read 192mm; number of slices 30; transversal orientation]. Three dummy scans were acquired at the beginning of each block. As well as the functional scans, an anatomical scan of the entire brain was acquired [3D MPRAGE; saggital; TE 2.37ms; TR 1800ms; flip angle 8o; voxel size 0.98mm × 0.98mm; FOV read 250mm; slice thickness 0.85mm; slices per slab 208; ascending acquisition; phase encoding direction A to P].

### fMRI Pre-processing

MRI images were pre-processed using the procedure recommended in SPM12 (Wellcome Department of Imaging Neuroscience, Institute of Neurology, London, UK). Functional images were first re-aligned per experimental block. These were registered to the mean image with a 6th degree spline interpolation. Following this was co-registration of the structural image to aligned functional images, segmentation of white and gray matter, normalisation of functional images using the deformation field created during segmentation, and normalisation of the functional to structural. The resulting data was smoothed with a 4-mm full-width-half-maximum Gaussian Kernel, and a high-pass filter with a cut off period of 128 s was applied. The registration of images was checked visually at each stage. Parameters not specified here can be assumed as the default SPM parameters.

### fMRI Analysis

During first level analysis, general linear models (GLM) with event-related designs were conducted in SPM12, to identify voxels activated in response to trial type (*blind*/*localise*/*sense*/*false alarm*/*correct rejection*). Regressors were created for each trial type by convolving the stimulus onset times with the canonical hemodynamic response function (HRF) across all blocks (Friston et al., 1994). Each regressor had a duration matched to the length of visual display, and serial correlations were corrected using the AR(1) method. All fMRI analysis was conducted in relation to the onset of the second display, where the change could occur. However, given the fast presentation of the two displays, it is possible that activation from the first display contributed to the activation recorded during the second. Each block was modelled with a separate set of regressors including time derivatives, as we did not perform slice time correction. Six motion regressors were added as nuisance variables.

For each participant we ran the following contrasts during first-level analysis; *sense > blind*, *localise > blind*, *localise > sense*, *blind* > no-change, *sense > false alarm*, *false alarm > sense*. We then compared awareness conditions at the second-level using one-sample t-tests. An additional paired-samples t-test was used to identify voxels with activation that was significantly different between the pair of contrasts *localise > blind* and *sense > blind*.

The contrasts *localise > blind* and *sense > blind* should reveal voxels with activation specific to full or partial awareness of the change, respectively, compared to no awareness. As these three conditions all contain a change in coloured square, the difference is the participant’s level of awareness. In the contrast *localise > sense*, we should identify voxels only activated when participants can both detect and localise the change, compared to only change detection. These areas would therefore be indicated in the facilitation of complete visual awareness, compared to *sensing* alone. We did not run the contrasts in the other direction, for example *blind > localise*, given previous results that suggest very little activation present for *blind* trials (Beck et al., 2001; Pessoa, 2004). Contrasting *blind* and no-change trials should reveal activation specific to the presence of the change, despite the participant being unable to detect it. The contrasts between *sense* and *false alarm* trials are useful to determine if *sensing* is similar to false alarms, meaning that participants did not detect anything changing during the change trials and were overconfident in their awareness.

To identify voxels with activation that correlated with the change in task difficulty over time, a separate GLM model was constructed with one regressor for the onsets of all trials, and a parametric regressor using the difficulty (or number of squares presented) at each trial. To identify voxels with activation that correlated with the change in participant certainty over time, a separate GLM model was constructed with one regressor for the onsets of all trials, and a parametric regressor using the certainty value reported by the participant at each trial.

Across all fMRI analyses, we report clusters with a minimum size of 20 voxels and a cluster-level family-wise error (FWE) corrected p < .001. Extended local maxima were labelled using two methods that provided overlapping results; the automated anatomical labeling (AAL) toolbox in SPM (12), with a local maximum radius of 5mm, and the SPM Anatomy toolbox, which for compatibility reasons used an older version of MATLAB (2014a) and SPM (SPM8). MNI co-ordinates were used to label voxels according to Brodmann areas. The SPM render function was used to plot our results on the cortex of an MNI brain. MRICron was used to create multi-slice views of the t-score maps for each contrast of interest.

### ERP-informed fMRI Analysis

For ERP-informed fMRI analysis, a first-level model with one regressor was constructed for the onset of all change trials (*blind/localise/sense*), with single-trial ERP values included as a parametric regressor. The LP ERP in response to the change display was chosen a-priori for this analysis, as significant differences have previously been identified between awareness conditions within this late parietal potential (Scrivener et al., 2019; Fernandez-Duque & Thornton, 2003; Busch et al., 2010). A second regressor was added for the onset of all no change trials. Motion parameters were also included as nuisance variables.

## Behavioural Results

### Accuracy and reaction times

Accuracy for question 1, in which participants had to identify a change, had a mean of 54% (range = 39 − 69%, *SD* = 9%). Accuracy for question 2, in which participants had to localise the change, had a mean of 72% (range = 61 − 86%, *SD* = 8%). The mean difficulty level given to each participant ranged from 6 to 23 squares (*M* = 16, *SD* = 4), with the maximum difficulty experienced by each participant ranging from 18 to 36 (*M* = 27, *SD* = 5). D’prime scores ranged from .940 to 2.30 (*M* = 1.38, *SD* = .38). One person had a negative criterion, meaning that they had a response bias towards false alarms. All other participants had positive criterion, indicating a conservative response strategy (*M* = .61, *SD* = .33). D’prime scores were significantly different from 0 in a one-sampled t-test, indicating that participants could discriminate between change and no change trials, *t*(19) = 16.263, *p* < .001.

Mean difficulty correlated with mean location accuracy (*r* = .543, *p* = .013) and d’prime (*r* = −.601*p* = .005), but not with mean detection accuracy (*r* = −.371, *p* = .107). Maximum difficulty also correlated with mean location accuracy (*r* = .537, *p* = .015) and d’prime (*r* = −.482, *p* = .031), but not with mean detection accuracy (*r* = −.349, *p* = .131).

The percentage of *false alarm* trials (12.23% ± 8.64) was lower than the percentage of *sense* trials (28.07% ± 7.73) *t*(19) = −6.815, *p* < .001, *g*_*rm*_ = 1.85, suggesting that *sense* trials occurred more often than participants made false alarms. Additionally, the percentage of false alarms was not significantly correlated with the percentage of *sense* trials (*r* = .198, *p* = .403).

Out of the 20 participants included in the analysis, 15 were slower to respond when they were *blind* to the change, compared to no-change trials. Reaction times for *blind* trials were also significantly slower than no-change trials (0.617 ± 0.176 s), *t*(19) = −3.613, *p* = .002, *g*_*rm*_ = 0.25. Therefore, despite being *blind* to the change, the presence of a change in the display increased reaction times, particularly for trials where the participant was uncertain.

We found a significant effect of location of the changed item (outside/central) on awareness (*blind*/*localise*/*sense*), *χ*^2^(2) = 26.68, *p* < ,001, as participants were more likely to be *blind* to the change when it occurred on the outside of the display (*blind* outside: 911 trials, central: 619). There were also a greater number of *sense* trials for outside changes, suggesting that these changes may be harder to localise than central changes (*sense* outside: 290 trials, central: 220). The location had the least influence on *localise* trials (*localise* outside: 627, central: 619).

The hemisphere of the display in which the change occurred (left/right) had no significant effect on participant awareness (*blind*/*localise*/*sense*), *χ*^2^(2) = 4.941, *p* = .085 (*blind* left: 781 trials, right: 749; *localise* left: 651, right: 607; *sense* left: 236, right 276). Additional behavioural analysis and results can be found in the supplementary material.

## EEG Results

### P1 and N1

For P1 amplitudes, the main effect of awareness was not significant, *F* (2, 38) = .568, *p* = .572., *η*^2^ = .029. Display was also not significant, *F* (1, 19) = .143, *p* = .709, *η*^2^ = .007. The interaction between awareness and display was not significant, *F* (2, 38) = 3.250, *p* = .050, *η*^2^ = .146 (figure 2). (*Blind* first display *M* = 1.933, *SD* = 4.106, second *M* = 1.401, *SD* = 5.052; *localise* first *M* = 0.606, *SD* = 2.706, second *M* = 1.108, *SD* = 5.858; *sense* first *M* = 0.509, *SD* = 2.738, second *M* = 2.020, *SD* = 5.900.)

**Figure 2.**
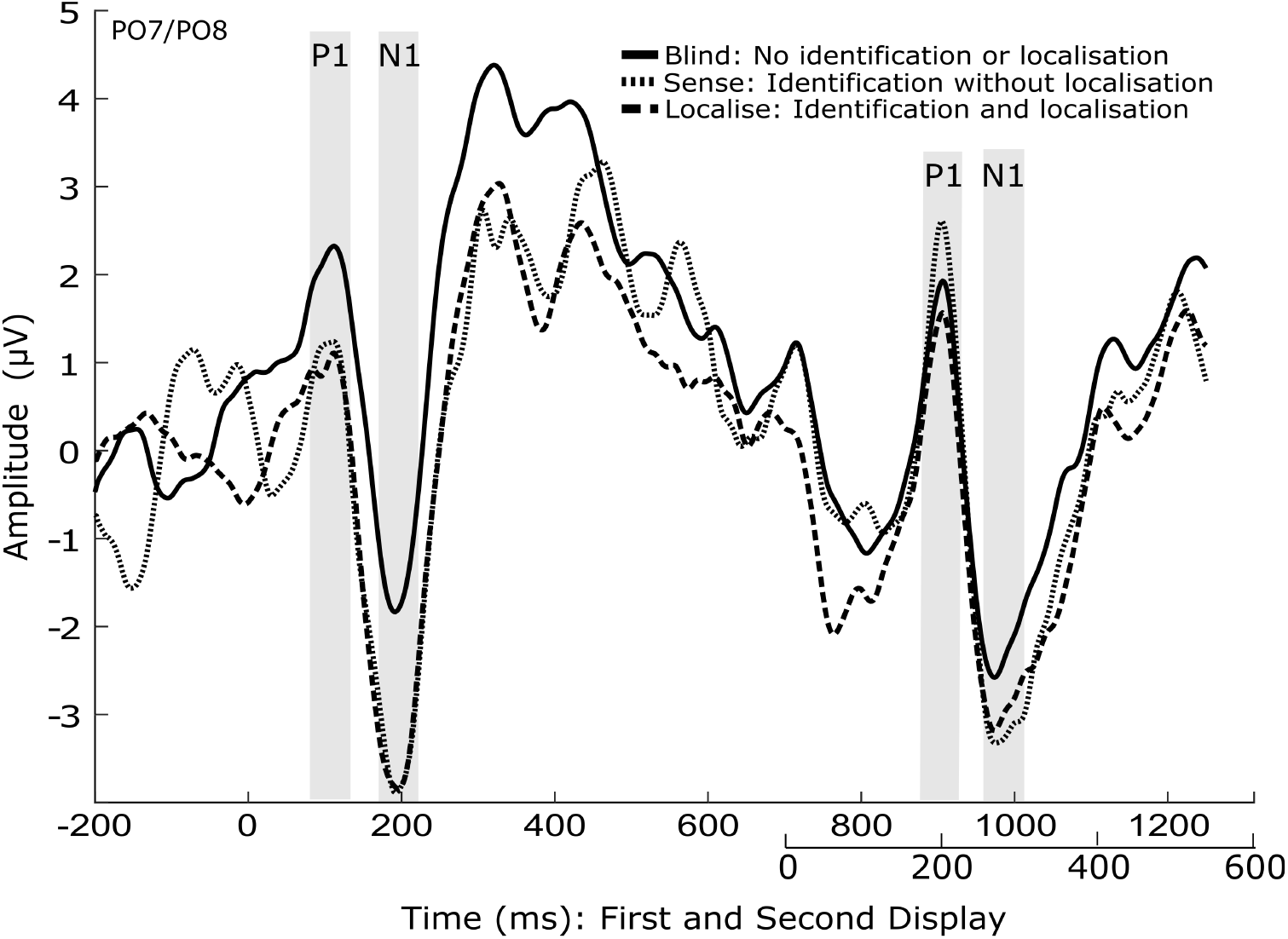
ERP plot showing the mean of electrodes PO7 and PO8, for each awareness condition. Condition means for the values within the shaded time windows were used for ERP analysis.

For the N1, the main effect of awareness was not significant, *F* (2, 38) = 2.008, *p* = .148, *η*^2^ = .096. Display was also not significant, *F* (1, 19) = .68., *p* = .797, *η*^2^ = .004, nor was the interaction between awareness and display, *F* (2, 38) = 2.046*p* = .143, *η*^2^ = .097 (figure 2). (*Blind* first display *M* = −1.526, *SD* = 4.096, second *M* = −2.178, *SD* = 4.469; *localise* first *M* = −3.609, *SD* = 4.246, second *M* = −2.783, *SD* = 5.658; *sense* first *M* = −3.500, *SD* = 3.662, second *M* = −2.881, *SD* = 5.279.)

### N2pc

The main effect of hemisphere on N2pc amplitudes was not significant, *F* (1, 19) = .338, *p* = .568, *η*^2^ = .018, nor was the main effect of awareness, *F* (2, 38) = .878, *p* = .424, *η*^2^ = .044. The interaction was not significant, *F* (2, 38) = .572, *p* = .569, *η*^2^ = .029.

As we had strong hypotheses about the presence of an N2pc for *localise* trials, we also ran corrected post-hoc pairwise comparisons across awareness levels. A significantly increased negativity across both hemispheres was found for *localise* trials (*M* = −1.573, *SD* = 4.378) compared to *blind* (*M* = −.810, *SD* = 4.856) *p* = .038. *Blind* and *sense* (*M* = −1.720, *SD* = 5.444) were not significantly different, *p* = .259, nor were *sense* and *localise*, *p* = .862.

### Visual Awareness Negativity (VAN)

The main effect of awareness on the VAN was not significant *F* (2, 38) = .029, *p* = .971, *η*^2^ = .002. (*Blind M* = 0.059, *SD* = 3.427, *localise M* = 0.184, *SD* = 3.093, *sense M* = 0.104, *SD* = 3.295.)

### Late Positivity (LP)

There was a main effect of awareness on LP amplitudes *F* (2, 38) = 3.776, *p* = .032, *η*^2^ = .166. In corrected post-hoc comparisons, *localise* trials (*M* = 2.270, *SD* = 4.130) had a significantly greater LP amplitude than *blind* (*M* = .032, *SD* = 2.158), *p* = .024. However, *sense* (*M* = 1.069, *SD* = 3.801) was not significantly different to *blind*, *p* = .130, or *localise* trials, *p* = .174 (figure 3).

**Figure 3.**
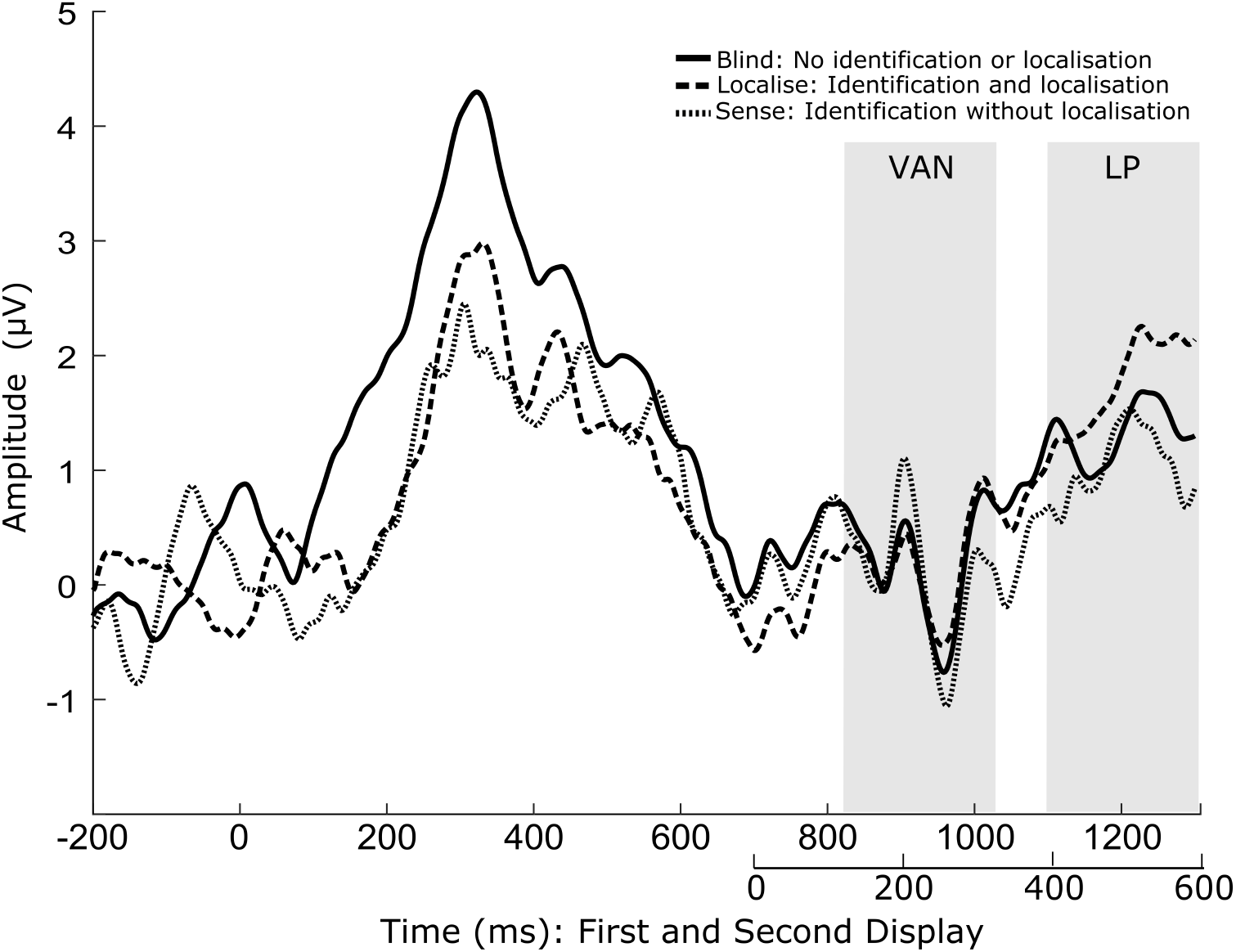
ERP plot showing a mean of electrodes Cz, CPz, and Pz, for each awareness condition. Condition means for the values within the shaded time window were used for ERP analysis.

## fMRI Results

### Awareness

For the contrast *localise>blind*, increased BOLD activation was found in the bi-lateral occipital cortex (BA17/V1, BA18/V2, and hOC4v/V4), bilateral parietal cortex (BA40/PFt, BA3b, BA2), left putamen (BA49), left fusiform gyrus (BA37), right insula (BA13), right pre-motor cortex (BA6, spanning middle frontal, superior frontal, and pre-central gyri), (see figure 4 for significant clusters, figure 5 for a map of t-scores, and table 1 for additional values).

**Table 1.** Voxels significantly activated for the contrast localise > blind, cluster FWE p < .001, minimum 20 voxels.

| Brain Region | BA | Hemisphere | Cluster size | MNI Coordinates |  |  | Z Score | T (peak level) | Cluster level FWE (p) |
| --- | --- | --- | --- | --- | --- | --- | --- | --- | --- |
|  |  |  |  | x | y | z |  |  |  |
| Lingual gyrus | 17 | Right | 12408 | 4 | -82 | -2 | 5.36 | 9.64 | <.001 |
|  | 18 (hOC4v) | Left |  | -22 | -70 | -6 |  |  |  |
|  | 18 | Left |  | -6 | -76 | 0 |  |  |  |
| Inferior parietal cortex | 40 (PFt) | Left | 421 | -38 | -30 | 42 | 4.71 | 7.33 | <.001 |
| Postcentral gyrus | 3b | Left |  | -44 | -30 | 52 |  |  |  |
|  | 2 | Left |  | -40 | -42 | 58 |  |  |  |
| Putamen | 49 | Left | 90 | -18 | 8 | 8 | 4.25 | 6.08 | <.001 |
|  |  |  |  | -22 | 0 | 6 |  |  |  |
| Middle frontal gyrus | 6 | Left | 182 | -28 | 4 | 56 | 4.18 | 5.89 | <.001 |
| Superior frontal gyrus | 6 | Left |  | -26 | -6 | 60 |  |  |  |
| Precentral gyrus | 6 | Left |  | -30 | -22 | 64 |  |  |  |
| Insula | 13 | Right | 96 | 28 | 22 | -10 | 4.08 | 5.66 | <.001 |
|  |  |  |  | 32 | 28 | -2 |  |  |  |
|  |  |  |  | 26 | 24 | 2 |  |  |  |
| Inferior temporal gyrus | 39 | Left | 84 | -56 | -56 | 12 | 3.71 | 4.87 | <.001 |
| Middle temporal gyrus | 37 | Left |  | -48 | -58 | -4 |  |  |  |
|  |  |  |  | -60 | -52 | -6 |  |  |  |
| Inferior parietal cortex | 40 (PFt) | Right | 110 | 44 | -24 | 38 | 3.62 | 4.69 | <.001 |
| Postcentral gyrus | 3b | Right |  | 50 | -18 | 40 |  |  |  |
|  |  |  |  | 40 | -28 | 46 |  |  |  |

**Figure 4.**
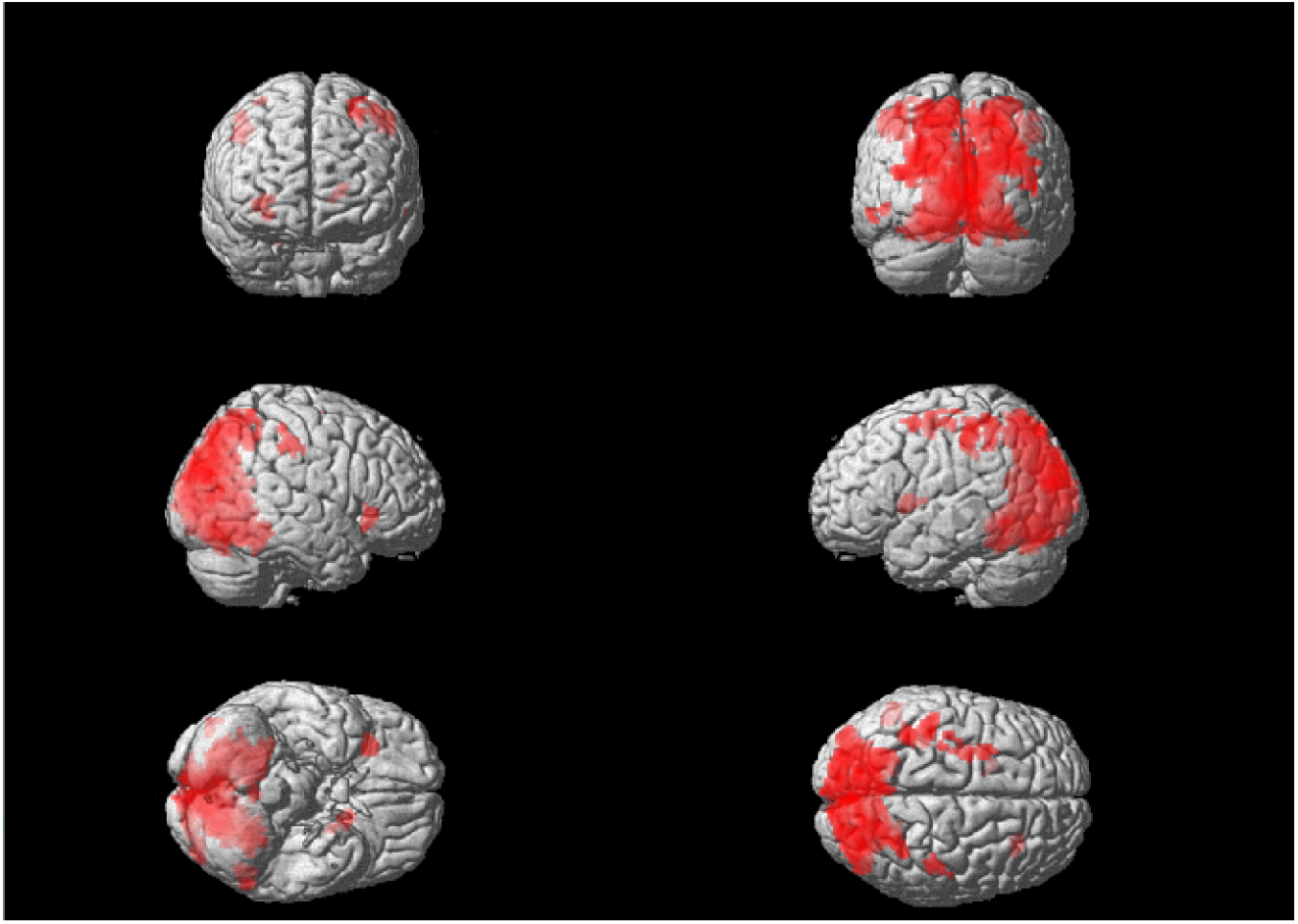
Voxels activated for the contrast localise > blind trials. Multiple comparisons were controlled using a cluster level family wise error correction where p < .001, as well as a minimum cluster size of 20 voxels.

**Figure 5.**
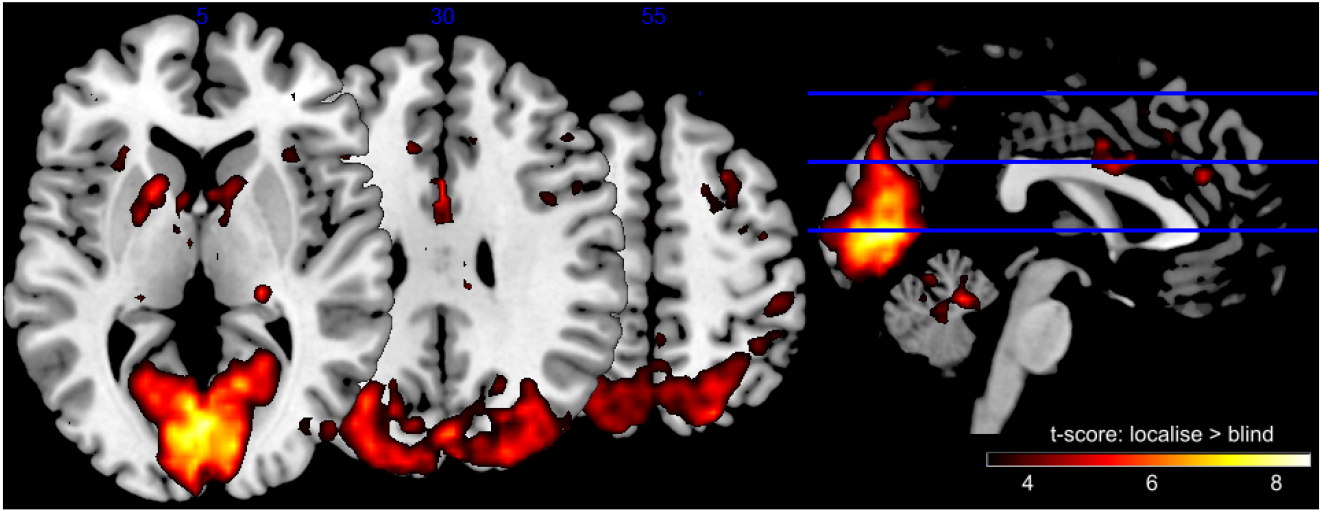
T-scores for the contrast localise > blind trials, thresholded at a minimum *t* = 3.

For the contrast *sense>blind*, increased activation was found in bilateral occipital cortex (BA17/V1, BA18/V2), left pre-motor cortex (BA6, spanning middle frontal, superior frontal, and precentral gyri), superior medial gyrus (BA8), parietal cortex (BA40, BA7/hIP3), and left anterior cingulate cortex (BA32), (see figure 6 for significant clusters, figure 7 for a map of t-scores, and table 2 for additional values).

**Table 2.** Voxels significantly activated for the contrast sense > blind, cluster FWE p < .001, minimum 20 voxels.

| Brain Region | BA | Hemisphere | Cluster size | MNI Coordinates |  |  | Z Score | T (peak level) | Cluster level FWE (p) |
| --- | --- | --- | --- | --- | --- | --- | --- | --- | --- |
|  |  |  |  | x | y | z |  |  |  |
| Precentral gyrus | 6 | Right | 145 | 32 | -4 | 48 | 4.94 | 8.07 | <.001 |
| Middle frontal gyrus | 6 | Right |  | 24 | 16 | 50 |  |  |  |
| Superior frontal gyrus | 6 | Right |  | 24 | 8 | 60 |  |  |  |
| Calcarine gyrus | 17 | Left | 5526 | -4 | -96 | 4 | 4.92 | 8 | <.001 |
|  | 18 | Right |  | 12 | -72 | 10 |  |  |  |
| Lingual gyrus | 17 | Left |  | -6 | -82 | 2 |  |  |  |
| Superior medial gyrus | 8 | Left | 88 | -2 | 26 | 0 | 3.96 | 5.38 | <.001 |
| Anterior cingulate gyrus | 24 | Left |  | -4 | 28 | 30 |  |  |  |
| Superior medial gyrus | 8 | Right |  | 8 | 24 | 42 |  |  |  |
| Inferior parietal sulcus | 40 | Left | 87 | -48 | -42 | 56 | 3.67 | 4.78 | <.001 |
|  | 7 (hIP3) | Left |  | -32 | -48 | 49 |  |  |  |
|  | 40 | Left |  | -40 | -44 | 50 |  |  |  |

**Table 3.** Parametric regressor: participant certainty, cluster FWE p < .001, minimum 20 voxels.

| Brain Region | BA | Hemisphere | Cluster size | MNI Coordinates |  |  | Z Score | T (peak level) | Cluster level FWE (p) |
| --- | --- | --- | --- | --- | --- | --- | --- | --- | --- |
|  |  |  |  | x | y | z |  |  |  |
| Lingual gyrus | 18 | Right | 160 | 14 | -74 | 0 | 4.11 | 5.74 | <.001 |
| Calcarine gyrus | 17 | Right |  | 6 | -82 | 8 |  |  |  |
|  |  |  |  | 10 | -88 | 12 |  |  |  |
| Inferior parietal cortex | 40 | Right | 104 | 50 | -34 | 48 | 3.84 | 5.12 | <.001 |
|  |  |  |  | 48 | -42 | 48 |  |  |  |
|  |  |  |  | 60 | -34 | 34 |  |  |  |

**Table 4.** Parametric regressor: task difficulty, cluster FWE p < .001, minimum 20 voxels.

| Brain Region | BA | Hemisphere | Cluster size | MNI Coordinates |  |  | Z Score | T (peak level) | Cluster level FWE (p) |
| --- | --- | --- | --- | --- | --- | --- | --- | --- | --- |
|  |  |  |  | x | y | z |  |  |  |
| Lingual gyrus | 18 | Left | 69 | -16 | -92 | -6 | 4.23 | 6.02 | 0.003 |
|  |  |  |  | -10 | -86 | -12 |  |  |  |

**Figure 6.**
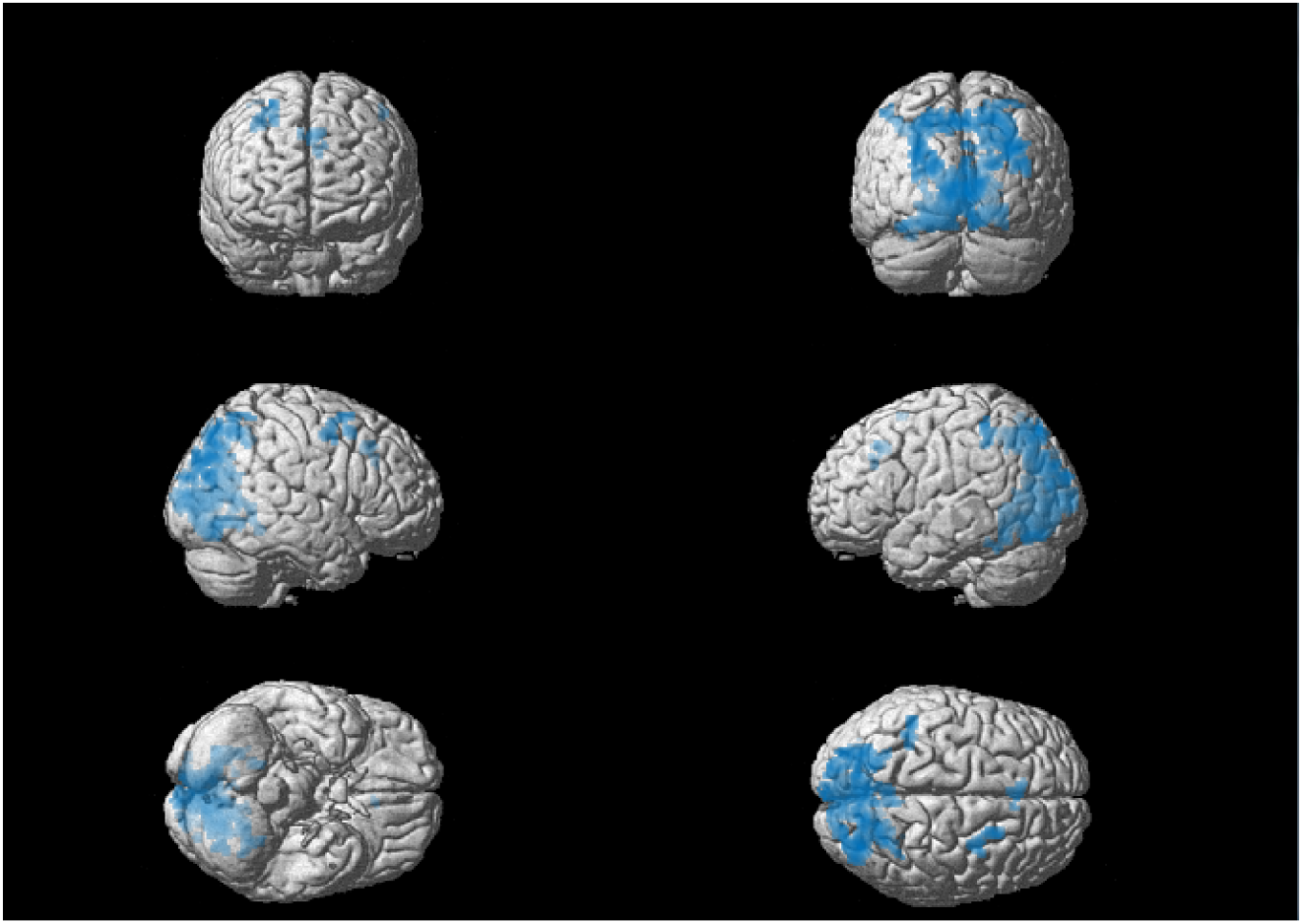
Voxels activated for the contrast sense > blind trials. Multiple comparisons were controlled using a cluster level family wise error correction where p < .001, as well as a minimum cluster size of 20 voxels.

**Figure 7.**
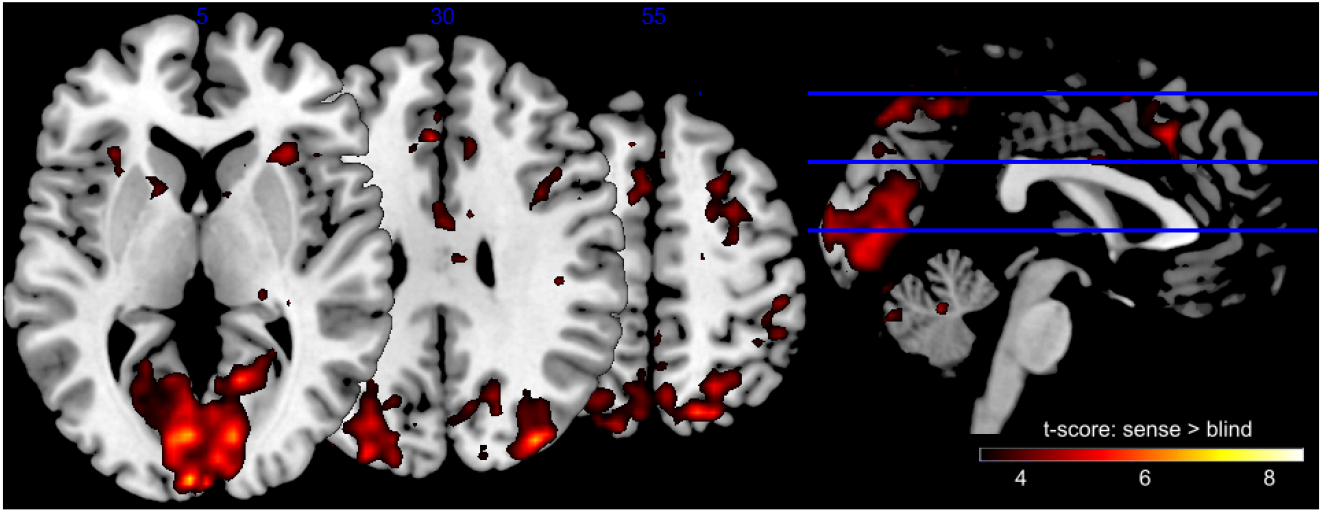
T-scores for the contrast sense > blind trials, thresholded at a minimum *t* = 3.

We also looked for any activation that was significantly greater in one contrast than the other (*localise>blind* vs. *sense>blind*). However, no significant activations remained after correction for multiple comparisons. No voxels survived for the following contrasts; *localise>sense*, *sense>localise*, *blind>*no change, *sense>false alarm*, or *false alarm>sense*.

### Post-hoc conjunction analysis

Given that the contrasts *localise* vs *blind* and *sense* vs *blind* revealed similar networks of activation, we ran a conjunction analysis to determine which voxels were significantly activated in both contrasts (note that this analysis was not included in our pre-registration). To do this, we entered the two first-level contrasts for each participant into an ANOVA at the second-level (independence not assumed). We then ran a conjunction analysis across both contrasts to identify common voxels, using the conjunction null hypothesis as suggested in Nichols et al. (2005). Significant activation was identified in the visual cortex (BA18, BA19) and inferior parietal cortex (BA7, BA39) (see table 5 and figure 8).

**Table 5.** Conjunction analysis: voxels activated for both localise versus blind and sense versus blind contrasts. Cluster FWE p < .001, minimum 20 voxels.

| Brain Region | BA | Hemisphere | Cluster size | MNI Coordinates |  |  | Z Score | T (peak level) | Cluster level FWE (p) |
| --- | --- | --- | --- | --- | --- | --- | --- | --- | --- |
|  |  |  |  | x | y | z |  |  |  |
| Lingual gyrus | 18 | Left | 6116 | -4 | -70 | 6 | 5.51 | 7.35 | <.001 |
|  |  |  |  | -4 | -80 | -8 |  |  |  |
|  |  |  |  | -18 | -72 | -10 |  |  |  |
| Inferior parietal cortex | 7 | Right | 445 | 28 | -76 | 44 | 4.91 | 6.16 | <.001 |
| Occipital cortex | 19 | Right |  | 22 | -90 | 22 |  |  |  |
| Inferior parietal cortex | 39 | Right |  | 34 | -76 | 38 |  |  |  |
| Inferior parietal cortex | 7 | Right | 421 | 18 | -70 | 42 | 4.73 | 5.84 | <.001 |
|  |  |  |  | 22 | -72 | 54 |  |  |  |
|  |  |  |  | 8 | -64 | 52 |  |  |  |

**Figure 8.**
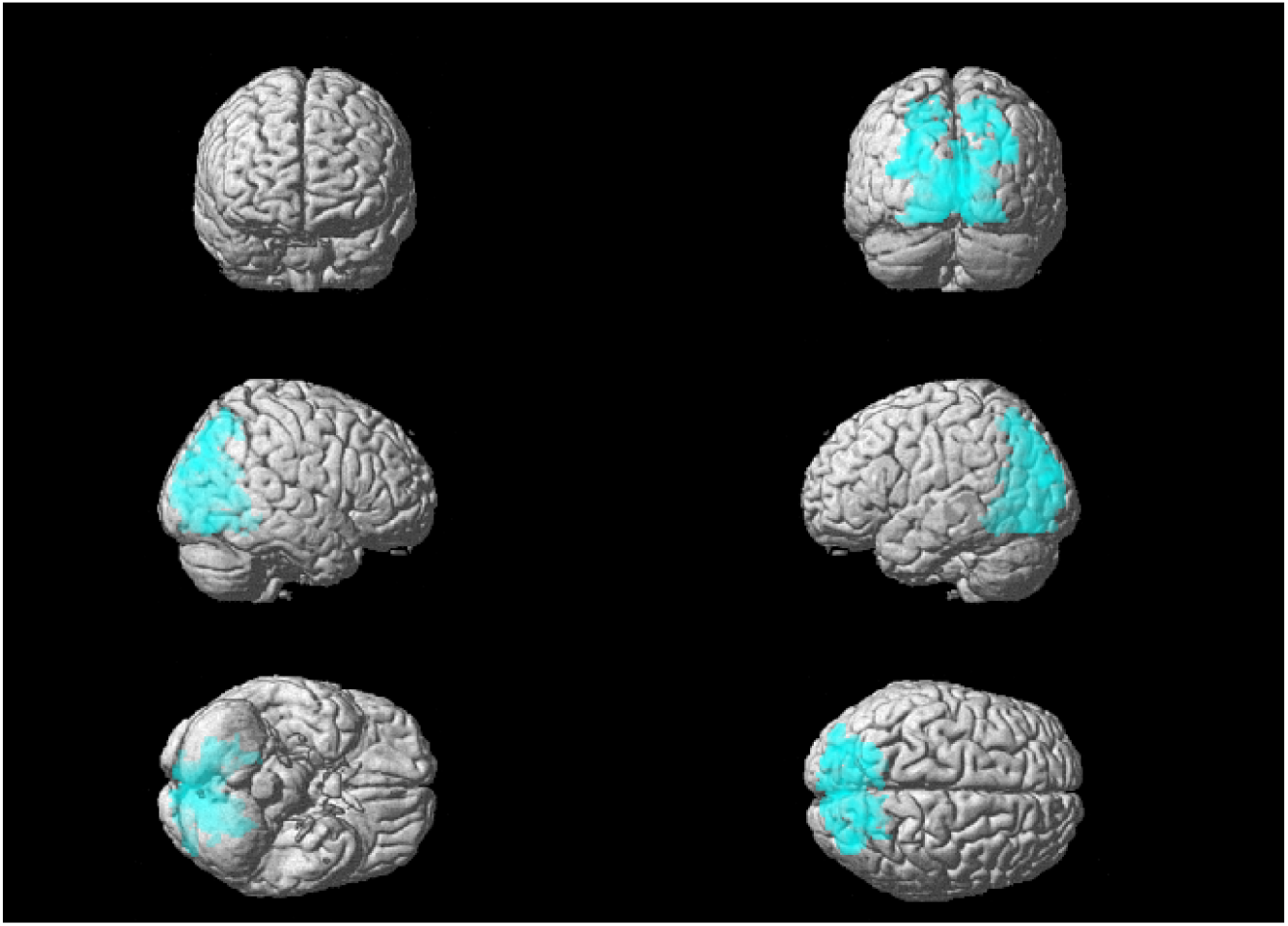
Conjunction analysis: voxels significantly activated for both *localise>blind* and *sense>blind* contrasts. Multiple comparisons were controlled using a cluster level family wise error correction where p < .001, as well as a minimum cluster size of 20 voxels.

### Difficulty and Certainty

The parametric regressor of participant certainty revealed significant activation in the right visual cortex (BA17/V1, BA18/V2) and right parietal cortex (BA40). The parametric regressor of task difficulty (the number of squares presented per trial) revealed significant activation in the left visual cortex (BA18, V2).

### ERP-informed fMRI

No significant voxels were identified for the LP-informed fMRI analysis using either method for extracting the single trial values.

## Discussion

The main aim of this change blindness experiment was to distinguish between trials in which participants could both detect and localise a change in coloured square (*localise*), versus those in which they could only detect it (*sense*), or not detect it at all (*blind*), using combined EEG-fMRI. In the EEG data, the late parietal positivity ERP, *localise* trials were significantly higher in amplitude than *blind* trials as previously found (Scrivener et al., 2019), but *sense* trials were not distinguishable from those where participants were *blind* to the change. Similarly, no differences were found between *sense* and *blind* trials in the N2pc or VAN. It is not clear whether this is due to false positive findings in the previous study, the smaller signal to noise ratio in the combined EEG-fMRI data, or the 530 relatively small sample size. The fMRI results revealed significant differences in BOLD activation for both *localise* and *sense* trials when compared to *blind*, suggesting that they are separable to trials where participants were completely unaware of the change. These results suggest that the *sense* condition may be distinguishable from the traditional *blind* condition, meaning that subjects may have access to more information when they are able to *sense* a change. However, the contrast between *localise* and *sense* conditions revealed no significant activations, and a conjunction analysis revealed overlapping activation in visual and parietal regions. These two levels of awareness may therefore be associated with activation within a similar network, and the link between brain activity and behavioural differences remains unclear.

### Behavioural

One explanation for the presence of a *sense* condition in change blindness is that it reflects a liberal response criteria, such that participants report seeing a change even though they were not certain that it occurred (Simons & Ambinder, 2005). In other words, they make a *‘false alarm’* during change trials. If this is the case, then these trials may be similar in number to *false alarm* trials, where participants incorrectly report a change for identical displays where they could not have seen a change. We found that participants had fewer *false alarms* than *sense* trial, and the percentage of these trials across participants was not correlated. This suggests that *sense* trials cannot be attributed to a liberal response criterion of the participants, as the tendency of participants to make a *false alarm* did not influence the number of times they could *sense* a change. However, this differs from previous results, where a significant correlation was found in the percentage of the two trial types (Scrivener et al., 2019). Further behavioural data may therefore be needed to confirm this relationship.

Previous studies have also reported that participants responded ‘no change’ more quickly for no-change trials, compared to change trials (Williams & Simons, 2000; Mitroff et al., 2004). The participant’s response is the same in both trial types, but the presence of a change is different. This suggests that even when they fail to detect the change in a change trial, they take longer to respond. We therefore compared reaction times for no-change trials and *blind* trials. Out of the 20 participants, 15 were slower to respond when they were blind to the change, compared to no-change trials. Reaction times for *blind* trials were also significantly slower than no-change trials, meaning that even when participants did not notice the change, its presence increased their reaction times. It is possible that in *blind* trials, some information may be available to the participant, leading to slower reaction times, but not enough for them to be confident to report the change.

The location of the square that changed in colour during the experiment had a significant influence on the likelihood that the change was detected; changes closer to the central fixation were detected at a higher frequency across participants than those further away. One explanation is that the participants were asked to fixate at the centre of the screen, and therefore their overt attention was directed here during the trial. As attention has been found to correlate with change detection, this finding is not surprising (Rensink et al., 1997).

### EEG

For the late parietal positivity ERP, *localise* trials were significantly higher in amplitude than *blind* trials. Other studies have also reported increased LP amplitudes for detected versus undetected changes (Fernandez-Duque & Thornton, 2003; Busch et al., 2010), which has been suggested to reflect conscious awareness of changes (Railo et al., 2011) and participant confidence (Eimer & Mazza, 2005). However, *sense* trials were not distinguishable from trials where participants were *blind* to the change. This contradicts our own results from a previous study where all three awareness conditions were distinguishable within the LP (Scrivener et al., 2019). There is therefore increasing evidence that the LP varies reliably between detected versus undetected changes, but whether it can be detected during *sense* trials is unclear. Note that the presence of a significant LP for *localise*, but not for *sense*, should not be used as evidence that the two are different as the post-hoc comparison was not statistically significant.

For the N2pc results, it should be emphasised that the main effect of hemisphere was not significant. Therefore, the post-hoc comparison in amplitude between *localise* and *blind* trials does not reflect the traditional asymmetry of the N2pc component, with a greater negativity in the contralateral hemisphere. It can only be concluded that there was an increased negativity for *localise* trials across both hemispheres, and may be better interpreted as an N2 component. This is a common finding, and in a review of the ERP correlates of visual awareness (Koivisto & Revonsuo, 2010) the majority of change blindness paper reported enhanced negativity in the N1-N2 range for detected changes (with the exception of Fernandez-Duque et al., 2003 and Neideggen et al., 2001).

In a previous EEG study we did find a significant N2pc for both *localise* and *sense* conditions, including a significant main effect of hemisphere (Scrivener et al., 2019). We concluded that the presence of an N2pc for both awareness conditions indicated a shift in attention towards the hemisphere of the change (Luck & Ford, 1998), but that this shift in attention was not sufficient to facilitate correct localisation in *sense* trials. In this experiment, we failed to find any evidence for this shift in either awareness condition, as characterised by the N2pc.

We found no statistically significant effects of awareness in the P1, N1, or VAN ERP analysis, similar to our previous results (Scrivener et al., 2019). In a recent review of the ERPs associated with visual awareness (Förster et al., 2020), the authors concluded that early P1 and N1 peaks are unlikely to be the earliest signature of visual awareness, and no longer discuss these peaks as possible candidates. This is due to increasing evidence against their association with conscious detection, which our findings support. However, they argue that the VAN is the most likely candidate for a marker of conscious detection, and our results are contrary to several previous findings. One possible explanation is the difference in experimental paradigm. In many cases, awareness is modulated by the perceptual difficulty of the stimuli, for example by the contrast. However, the stimuli in a change blindness paradigm remain at the same contrast across all trials, and difficulty is instead modulated by the number of distractors. Another suggestion from our previous work (Scrivener et al., 2019) is that the VAN requires both the location and identity of an object to be stored, such that it is available for conscious report. As our participants were not able to identify the location of the change in the *sense* condition, this may explain the lack of significant VAN ERP.

### fMRI

#### Awareness

One aim of this experiment was to improve our knowledge of the neurological basis of the *sense* condition with the addition of fMRI results. We found largely overlapping activation for both *localise* and *sense* conditions when contrasted with trials where participants were *blind* to the change in coloured square. Both awareness conditions had significantly greater activation in the early visual cortex (B18, V2), the left supramarginal gyrus in the inferior parietal lobe (BA40), and the left pre-motor cortex (BA6).

The posterior parietal cortex and early visual cortex are commonly implicated as storage sites for the contents of visual working memory (Todd and Marois, 2004; Edin 2009; D’Esposito 2015), and previous fMRI studies of change detection also found activations in these areas (Beck et al., 2001; Pessoa, 2004). Using MVPA, Christophel et al (2012) identified stimuli-specific information contained in both early visual and posterior parietal areas (around the intraparietal sulcus), further implicating these regions as storage sites for visual representations. The activation of these visual and parietal regions in both *localise* and *sense* conditions suggests the presence of visual representations of the stimuli for both levels of awareness. This supports the hypothesis that change blindness may arise from a failure to compare two displays or images, rather than a failure to encode the visual information (Simons et al., 2005; Hollingworth et al., 2001). Therefore, the inability of participants to localise the change during *sense* trials may not be explained by a lack of parietal representation, as activity in the dorsal stream (BA18 and BA40) was greater than during *blind* trials.

Activation found only in the *localise* contrast (but not for *sense*) were located in the primary sensory cortex (BA2, BA3b), putamen (BA49), and insula (BA13). This forms a wider network of activation than the *sense* versus *blind* contrast, including mid-brain structures. The insula and putamen are both hypothesised to act as hubs in key brain networks relating to cognitive control, and their activation specific to *localise* trials may indicate their role in facilitating full awareness of the change. More specifically, the insula forms an integrative hub between attention and salience networks (Menon & Uddin, 2010; Eckert et al., 2009), balancing external attentional cues with internal performance monitoring (Uddin et al., 2017). In contrast, the putamen is suggested to be a central component of a frontal-subcortical network (including the superior parietal and premotor cortex) related to cognitive control (van Belle et al., 2014), and has anatomical connections with rostral parietal areas (Jarbo & Verstynen, 2015). Further, patients with putamen lesions show symptoms of left-sided neglect (Karnath & Rorden, 2012), which is often thought of as a disorder of attention.

Overall, the pattern of findings indicates both anatomical and functional links between the putamen/insula and parietal cortex, which may explain their increased activation during *localise* trials. However, it should be noted that our fMRI sequence parameters were not specifically designed for accurate recording of mid-brain structures, which may influence the reliability of these results (Eapen et al., 2011).

Activation in the anterior cingulate cortex (ACC) was found in the *sense* versus *blind* contrast. The ACC is commonly linked to functional networks underlying attention (Ungerleider, 2000), and more specifically in boosting attention towards task-relevant stimuli (Orr & Weissman, 2009; Kim et al., 2016). Further, Mitchell and Cusack (2008) found ACC activation that correlated with estimates of the number of items stored by each participant during a working memory task. If this activation reflects increased attention towards the changed stimuli, then it would be expected to occur in both awareness conditions, as attention facilitates change detection (Rensink et al., 1997). However, ACC activation was not found in the *localise* condition, and therefore may not be necessary for full awareness of the change.

A more fitting explanation of the ACC activation specific to the the *sense* condition is that it reflects error processing during the task. This is because *sense* trials contained a response error, where participants incorrectly localised the change. Using combined EEG-fMRI, ACC activation has been linked to error processing and is correlated with the error related negativity (ERN) in EEG (Iannaccone et al., 2015; Debener, 2005). Activity in this area could therefore relate to the incorrect responses of the participants during *sense* trials. However, it should be noted that activation in the ACC is found for a wide range of tasks and the specificity of this activation is debated (Dehaene, 2018).

It could be argued that *blind* trials also contain a response error, as the participant failed to report a change that did occur. This should therefore also activate the ACC, if ACC activation reflects error monitoring (and that this error monitoring need not be conscious). Compared to *blind* trials, *sense* trials contained activation in visual (BA18) and parietal (BA40) areas, and the participant correctly reported the change. However, it is also possible that the ACC activation relates to the participant’s awareness of their own failure to localise the change, which is not relevant to *blind* trials where the participant can be very confident that no change occurred. Further, the ACC activation during *sense* trials could reflect a mismatch between the intended response and the actual response (Dehaene, 2018). Although participants had represented the stimuli in visual working memory (indexed by the increased visual and parietal activation that was similar to *localise* trials), and planned the correct response, their actual response did not match their intended one leading to ACC activation. In *blind* trials, participants had significantly reduced visual and parietal activation, and may not have known which response was correct. Therefore, this mismatch between intended correct response and actual response did not occur. While this may explain our results, this is currently a working theory that should be explored in further research.

In relation to theories of visual consciousness, our results could be interpreted in support for the ‘partial awareness hypothesis’ given the distinction in fMRI between *blind* and *sense* trials. Although participants were aware of the change during *sense* trials, their inability to provide further information suggests a less detailed representation of the visual display. Further, *localise* trials were associated with similar activity to *sense* in visual and parietal areas, perhaps reflecting activity relating to the ‘all-or-nothing’ ignition of change detection. However, the additional activation related to *localise* trials may characterise an improved representation that facilitated correct localisation. This hypothesis is highly speculative, and clarity is needed on the distinction between *localise* and *sense* conditions. For example, with future work using MVPA it would be possible to determine if the pattern of information stored within the brain is similar between these two levels of awareness. This would provide more information regarding the nature of stored representations during the task, and identify regions where these representations differ. Given the behavioural and phenomenological differences between *localise* and *sense* trials, it is reasonable to expect that somewhere in the brain should contain differing representations for these two levels of awareness, and therefore be driving the variation in participant response.

#### Difficulty and certainty

Using participant certainty at each trial as a parametric regressor, we found significant activations in the right visual cortex (BA18, V2) and bilateral supramarginal gyrus (BA40). These regions were also found to increase with awareness of the change (*localise* and *sense* trials), possibly due to the relationship between awareness and certainty. Specifically, when participants were aware of the change and could localise it correctly, they were likely to report higher certainty in their responses.

The parametric regressor of task difficulty (the number of squares presented per trial) revealed significant activation in the visual cortex (BA18, V2). This finding likely reflects the greater visual stimulation associated with a more complex visual array. In previous literature, parietal activity has also been correlated with set size and the number of objects stored in visual working memory (Mitchell & Cusack, 2008). Activity also predicts individual differences in working memory capacity (Vogel & Machizawa, 2004). We failed to find this effect, which may be explained by the variation in set sizes that were presented across participants. Instead of presenting a number of blocks with a number of difficulty levels, the difficulty was modulated in real time depending on participant performance. Also, the change in response may not be linear in our case; during easy trials, the response may scale linearly with the number of trials, until the maximum capacity of the participant is reached. Past this point, the number of items may exceed the capacity, and therefore fail to be represented or modulate the brain activation in these regions.

### ERP-informed fMRI

Our pre-registered analysis method of LP-informed fMRI revealed no significant results. We therefore failed to identify voxels with activation that significantly co-varied with fluctuations in the EEG. It is acknowledged that EEG-BOLD couplings are weak, as they measure the effects remaining after the mean evoked BOLD responses are explained (Liu et al., 2016). However, previous combined EEG-fMRI experiments have managed to identify correlates of EEG using ERP-informed fMRI (Debener, 2005; Eimer & Mazza, 2005), even if at liberal correction thresholds.

One possible reason for the failure to find significant ERP-informed BOLD effects is the reduced signal to noise in EEG signals recorded inside the MRI environment. A second possibility is the method that we used to quantify single-trial ERPs. There is no single method for ERP-informed fMRI analysis, and we therefore chose to run two separate analysis pipelines in case of disparaging results. In the first, we used raw values from the EEG time series. This method is susceptible to noise artifacts, and any trials in which the noise signal is greater than the neurological signal of interest will reduce the chance of observing an effect across conditions. Given the increased number of artifacts in EEG-fMRI data, and the absence of perfect artifact removal routines, it is possible that the signal to noise ratio was too small in the raw single-trial ERP values.

In the second method, we used ICA to identify components matching our ERP of interest, with the hypothesis that noise signals would have a reduced contribution to the single trial values extracted from this component (Debener, 2005; Wirsich et al., 2014). However, this method also produced null results in ERP-informed fMRI analysis. A downside to this method is that its success is dependent on a) the algorithm accurately separating independent components, and b) the correct selection of the components containing the ERP of interest. Other possible processing steps used in ERP-informed fMRI include linear classifiers (Walz et al., 2015; Goldman et al., 2009), autoregressive models (Nguyen et al., 2014), and spatial laplacian filters (Liu et al., 2016), to name only a few. However, it is not within the scope of our pre-registered analysis to adjust the pre-processing or analysis steps any further.

### Conclusions

Overall, one of the main aims of this experiment was to establish if the *sense* condition is separable from other awareness conditions in neural signals, as measured using EEG and fMRI. While the phenomenological experience of *sensing* differs from full awareness, it remains unclear whether this arises from a distinct state of neural activation, or whether these trials can be explained by explicit behavioural mechanisms such as participant response errors or lack of confidence. The strongest evidence presented here is the difference in fMRI activation for *blind* trials compared to *sense* trials. Across our sample, there was a greater spread of activation within areas such as the early visual cortex and inferior parietal sulcus when participants suspected a change, compared to when they missed it completely. This suggests that *sense* trials were measurably different to *blind* trials, and that participants did have access to more information regarding the change.

However, the contrast between *sense* and *localise* trials, where participants had full awareness, revealed no significant differences in activation. Additionally, the conjunction analysis revealed an overlapping occipitoparietal network of activation for these two levels of awareness. This suggests common activity related to the awareness of the change itself. In line with the ‘partial awareness hypothesis’, it may be that a degraded representation of the visual display within these regions contributed to failed localisation during *sense* trials.

While we attempted to distinguish between true *sense* trials and *localise* trials with an error using participant certainty, the number of *sense certain* responses was low. This meant that dividing the awareness conditions into certain/uncertain for EEG or fMRI analysis was not feasible. Future experiments could focus on obtaining higher trial numbers, which would hopefully facilitate this analysis. However, the very nature of the *sense* condition means that participants are unlikely to be ‘certain’ during many of the trials. One way around this would be to include a response option for participants to indicate if they think that they made a response error, although this would only identify trials where the participants were aware of their mistake.

In summary, our data suggests that the phenomenological experience of *sensing* a change is associated with increased activity in visual, parietal, and anterior circulate cortices, when compared to change blind trials. Given this increased activation including areas that are commonly implicated as the storage sites of visual working memory, we argue that *sensing* may not be caused by a lack of representation of the visual display. Instead, *sensing* may reflect unsuccessful comparison of the two displays (Simons et al., 2005; Hollingworth et al., 2001), or a degraded representation that prevents accurate localisation of the change in space.

## Acknowledgements

Thank you to Arran Reader, David Scrivener, and Zola Dean for their valuable comments on the manuscript, and to Isil Bilgin and Sophie Szymkowiak for their help with data collection. Thank you also to Maximilian Zangs for his help with the initial version of the experimental paradigm script. This research was funded by the EPSRC, EP/1503705 DTG 2014/2015.

## Conflict of Interest Statement

The authors declare that the research was conducted in the absence of any commercial or financial relationships that could be construed as a potential conflict of interest.

## Data Accessibility Statement

The raw data, pre-processed data, and analysis scripts can be found on the Open Science Framework: https://doi.org/10.17605/OSF.IO/W6BH3

## Notes

https://osf.io/w6bh3/

### Competing Interest Statement

The authors have declared no competing interest.

### Summary of Updates

Revision to correct a LaTeX compile error for figure 4.

