## Supplementary analysis and results for "Sensing and seeing associated with overlapping occipitoparietal activation in simultaneous EEG-fMRI" (from another version of this preprint: version 1, 2020-07-09)

### Supplementary analysis and results: sensing and seeing associated with overlapping visuoparietal activation in simultaneous EEG-fMRI

#### Reaction times

Reaction times were compared using a repeated measures ANOVA calculated for question (1/2/3) and accuracy (correct/incorrect). The main effect of question was not significant, $F(2, 38) = 3.077, p = .058$. (Q1: $M = .585$, Q2: $M = .578$, Q3: $M = .652$). There was a significant main effect of accuracy, $F(1, 18) = 43,899, p < .001$, as reaction times for correct answers were faster ($M = .568$) than incorrect answers ($M = .642$). The interaction between question and accuracy was also significant, $F(2, 38) = 3.857, p = .030$. This was driven by the fact that accuracy did not significantly modulate responses for Q2 (correct $M = .567$, incorrect $M = .588$). However, for Q1 and Q3, incorrect answers were slower than correct answers (Q1 correct $M = .535$, incorrect $M = .636$, Q3 correct $M = .602$, incorrect $M = .702$). These findings suggest that incorrect answers were slower than correct answers, but only for Q1 and Q3. The certainty question (Q3) had the largest reaction times overall.

#### Accuracy

The percentage of trials across conditions was compared with a repeated measures ANOVA for accuracy (correct/incorrect localisation) and certainty (certain/uncertain). There was a significant main effect of accuracy, $F(1, 19) = 6.788, p = .017$, with a higher percentage of correct ($M = 49.924$) than incorrect trials ($M = 48.779$). The main effect of certainty was also significant, $F(1, 19) = 12.720, p = .002$, with a higher percentage of certain trials ($M = 62.873$) than uncertain ($M = 35.830$). The interaction between accuracy and certainty was also significant, $F(1, 19) = 52.836, p < .001$. This was driven by the similar percentage of certain ($M = 48.599$) and uncertain trials ($M = 48.960$) when participants incorrectly localised the change. Conversely, when participants correctly localise the change, they had more certain ($M = 77.148$) than uncertain trials ($M = 22.700$). Participants were more certain of their responses when they were able to localise the change (*localise* trials). When they were unable to localise the change (*sense* trials), they were equal in their responses of certain and uncertain.

#### Difficulty

As shown in figure 1, a range of difficulty distributions were demonstrated by the participants, supporting our hypothesis. The averaged difficulty distribution across all participants can be found in figure 1. The mean difficulty level given to each participant ranged from 6 to 23 ($M = 16, SD = 4$), with the maximum difficulty experienced by each participant ranging from 18 to 36 ($M = 27, SD = 5$). Mean difficulty correlated with mean location accuracy ($r = .590, p = .008$) and d'prime ($r = -.601p = .005$), but not with mean detection accuracy ($r = -.371, p = .107$). Maximum difficulty also correlated with mean location accuracy ($r = .537, p = .015$) and d'prime ($r = -.482p = .031$), but not with mean detection accuracy ($r = -.349, p = .131$).

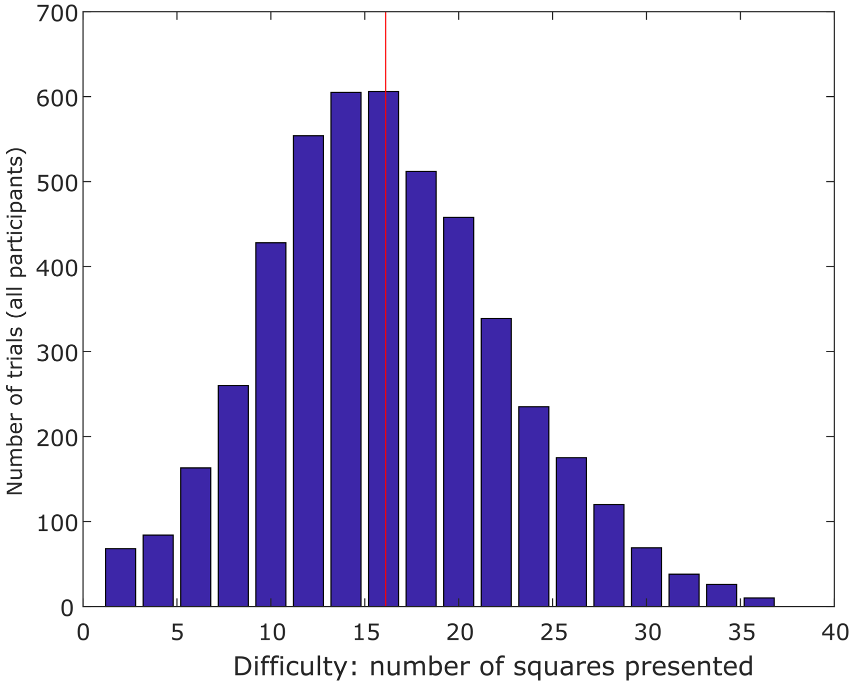


*Figure 1*. Histogram of task difficulty across the group. The red line represents the mean difficulty.

#### Colour

A chi-square test was used to compare the frequency of each colour change occurring across awareness conditions (*see*/*blind*). Due to the large number of possible random colour changes, *localise* and *sense* trials were combined into *see* trials. Each possible colour change was coded with a number between 1 - 21 (such that a change in either direction was codes with the same number). Colour changes with an adjusted standardised residual greater than 1.96 were considered to have a significant difference between conditions.

There was a significant effect of colour change on awareness, $\chi^2(20) = 120.342, p < .001$. Colour change pairs with a significantly higher number of *blind* trials were: yellow/green, yellow/cyan, green/cyan, blue/cyan, white/cyan. Colour change pairs with a significantly higher number of *see* trials were: magenta/red, cyan/red, blue/red, white/red.

The main finding from these results is that colour changes including red were much more likely to be detected, given the prevalence of colour pairings that included red in the *see* condition. The other findings is that colour changes including cyan were more likely to be missed, as there were more *blind* trials with cyan in the colour pairing.

#### Change location

During the localisation question, participants were asked to identify the location of the change square using a 2 x 2 response grid. The display was therefore divided into four squares: upper left, lower left, upper right, and lower right. During *sense* trials, participants reported an incorrect location for the change. The purpose of this analysis was to determine if, for incorrect answers, participants were more likely to identify the correct side of the display (left/right) or the correct field (upper/lower).

Nine out of twenty participants had more trials where they correctly identified the correct side of the display (left/right), but the incorrect hemisphere (upper/lower). The remaining eleven had more trials where they were more likely to report the correct hemisphere (upper/lower) but incorrect side (left/right). In an paired t-test, we found no significant difference between the number of trials in each condition, $t(19) = -0.697, p = 0.494$. There is therefore no clear distinction between the likelihood of identifying the correct side or hemisphere for incorrectly localise trials.

#### Single-trial ERPs

Single-trial ERP values for each participant, for each ERP, at each channel, were correlated with the difficulty and certainty levels across trials. After correcting for multiple comparisons using FDR correction ($q = .05$), no significant correlations were found for either task difficulty or participant certainty.
